# Transmembrane protein 97 is a potential synaptic amyloid beta receptor in human Alzheimer’s disease

**DOI:** 10.1101/2021.02.01.428238

**Authors:** Martí Colom-Cadena, Jamie Toombs, Elizabeth Simzer, Jane Tulloch, Rosemary J Jackson, James H Catterson, Jamie Rose, Lora Waybright, Anthony O Caggiano, Caitlin Davies, Monique Hooley, Sophie Dunnett, Robert Tempelaar, Makis Tzioras, Mary E Hamby, Nicholas J Izzo, Susan M. Catalano, Colin Smith, Owen Dando, Tara L. Spires-Jones

## Abstract

Synapse loss correlates with cognitive decline in Alzheimer’s disease, and soluble oligomeric amyloid beta is implicated in synaptic dysfunction and loss. An important knowledge gap is the lack of understanding of how amyloid beta leads to synapse degeneration. In particular, there has been difficulty in determining whether there is a synaptic receptor that binds amyloid beta and mediates toxicity. While many candidates have been observed in model systems, their relevance to human AD brain remains unknown. This is in part due to methodological limitations preventing visualization of amyloid beta binding at individual synapses. To overcome this limitation, we combined two high resolution microscopy techniques: array tomography and Förster resonance energy transfer (FRET) to image over 1 million individual synaptic terminals in temporal cortex from AD (n=9) and age matched control cases (n=6). Within postsynaptic densities, amyloid beta generates a FRET signal with transmembrane protein 97, cellular prion protein, and postsynaptic density 95. Transmembrane protein 97 is also present in a higher proportion of postsynapses in Alzheimer’s brain compared to controls. Further, we inhibited amyloid beta / transmembrane protein 97 interaction in a mouse model of amyloidopathy by treating with the an allosteric modulator CT1812 or vehicle. CT1812 drug concentration correlated negatively with synaptic FRET signal between transmembrane protein 97 and amyloid beta. In human induced pluripotent stem cell derived neurons challenged with human Alzheimer’s brain homogenate, transmembrane protein 97 and amyloid beta are present in synapses. Transcriptional changes are induced by Aβ including changes in genes involved in neurodegeneration and neuroinflammation. CT1812 treatment of these neurons caused changes in gene sets involved in synaptic function. These data support a role for transmembrane protein 97 in the synaptic binding of amyloid beta in human Alzheimer’s disease brain where it may mediate synaptotoxicity.

## Introduction

In Alzheimer’s disease, synapse loss is an early event in the aetiology of the disease and is the strongest pathological correlate of cognitive decline.[16, 56, 65] The mechanism(s) underlying synapse degeneration, however, are still largely unknown.[10] We and others have observed that oligomeric amyloid beta (Aβ) peptide causes synaptic dysfunction, accumulates within in synapses, and is associated with synapse loss around plaques.[34, 41, 52, 60, 64] While it is clear that toxicity of tau and changes in non-neuronal cells are also important in disease pathogenesis,[23] substantial evidence supports an important role for Aβ in synaptotoxicity early in the disease process in Alzheimer’s.[38] As such, it is important to identify synaptic binding partners of Aβ which may mediate synaptotoxicity in human brain. Disrupting binding of Aβ with synaptic receptors is a promising therapeutic avenue as such interactions are “druggable”, or able to be interrupted with standard pharmacological approaches.[13]

Synaptic Aβ binding partners have been identified in cell culture systems and mouse models, but their human relevance is still debated (reviewed in [5, 30, 46, 63]). Among the Aβ binding candidates, cellular prion protein (PrP_c_) represents the most studied, either alone or through a complex with mGluR5.[37, 62, 68, 75] Other suggested Aβ binding partners at synapses include the α7-nicotinic receptor,[49] Ephrin A4,[69] PSD95[35, 50, 52] and LilrB2.[32] An important outstanding question in the field is which of these potential partners binds Aβ in human synapses, as most binding partners have not been validated in Alzheimer’s cases nor using human derived Aβ species.[38, 63] Further, in model systems, Aβ is often overexpressed or applied exogenously, and due to the “sticky” nature of Aβ oligomers, this can result in false positive signals for interacting partners, which has been highlighted as a problem for translation in the field.[5]

TMEM97, transmembrane protein 97, is a promising potential synaptic binding partner of Aβ. TMEM97 was recently identified as the gene that codes for the Sigma-2 receptor.[3] Sigma-2 receptors have been studied for more than four decades and are drug targets for several conditions including cancer, pain and diverse CNS disorders;[21, 58] and most notably, a Sigma-2 modulator, CT1812, is in clinical development in Phase 2 trials for Alzheimer’s disease and dementia with Lewy bodies.[11, 20] In the context of Alzheimer’s, in 2014, Izzo and colleagues found that Sigma-2 modulators including CT1812 could displace Aβ synthetic oligomers from their synaptic receptors in cellular models and could improve cognitive deficits in a mouse model of Alzheimer’s.[25] Yi *et al* and Mondal *et al* similarly found that Sigma-2 receptor ligands prevent neurodegeneration in a worm model expressing human amyloid precursor protein.[44, 73]

Little is known about the pathophysiological role of Sigma-2, especially due to its unknown identity until the identification of TMEM97. TMEM97, initially known as MAC30,[47] is overexpressed in some cancers and it is believed to be a key player of cholesterol homeostasis[74] and calcium regulation.[8, 70, 73] Linking this function to Alzheimer’s, in cellular models, TMEM97 has recently been shown to form a ternary complex with Progesterone receptor membrane component 1 (PGRMC1) and LDLR[55] that may control the internalization of monomers and oligomers of Aβ.[54] In addition, our group recently found increased levels of TMEM97 in synaptoneurosomes from Alzheimer’s cases, compared to control, in a proteomics study,[24] supporting a potential role in synaptotoxicity in humans.

Until the 2017 discovery that *TMEM97* encodes the sigma-2 receptor,[3] the understanding of the ability of sigma-2 receptor modulators to displace Aβ oligomers from synapses was based off solely pharmacological/functional data, and data indicating that this effect was correlated with PGRCM1 expression. To date, there has been no direct interrogation of whether TMEM97, PGRMC1 and Aβ are found within the same synapses in human brain, whether CT1812 affects TMEM97 binding to Aβ within synapses, or whether PrPC binding to TMEM97 may underlie the ability of sigma-2 receptor modulators to displace Aβ oligomers. The herein paper addresses these knowledge gaps for the first time.

The study of synapses in the human brain represents a technical challenge due to their small volumes, which are smaller than the diffraction limit of light microscopy, making colocalization studies difficult. In the present work we applied a new approach for the study of the close proximity of proteins in synapses in human postmortem brain tissue. To visualize the potential interaction between Aβ and potential binding partners at synapses, we combined array tomography[43] and Föster resonance energy transfer (FRET) microscopy.[19, 76] Array tomography allows us to reach a 70nm axial resolution, which enables the identification of single synaptic terminals in three dimensions.[31] The combination of array tomography with FRET enhances the lateral resolution to ∼10nm in the selected single synaptic terminals allowing us to determine whether proteins are close enough to be interacting.[22, 52]

The current study tests the hypothesis that TMEM97 interacts with Aβ in synapses in human Alzheimer’s brain and that modulating this can recover synaptic phenotypes in model systems. We demonstrate that TMEM97 is a potential Aβ synaptic binding partner in human brain tissue and confirms that Sigma-2 receptor complex allosteric modulator CT1812 can reduce interactions between TMEM97 and Aβ *in vivo*. These findings shed light on the mechanisms of action by which a Sigma-2 receptor modulator may be acting in the context of Alzheimer’s, and may help further Alzheimer’s therapeutic approaches, in both drug discovery and clinical development. Finally, this study enables further technical advances in the study of the still elusive synaptic structures involved in neurodegeneration.

## Materials and methods

### Human cases

Patients fulfilling clinical and neuropathological criteria for Alzheimer’s disease (n = 9),[45] or cognitively healthy control cases (n = 6) were included in this study. Sample sizes were based on power calculations using effect size of 0.79 from our previous human array tomography studies looking at colocalization of clusterin and Aβ in synapses synapses in Alzheimer’s[29] indicating that n=6 per group is sufficient at power =0.8 to detect a difference between colocalization of proteins at synapses between AD and controls (calculated using the WebPower package in R 4.1.2). A post-hoc power calculation using the results from the primary question in this study – whether Aβ and TMEM97 generate a FRET signal in human synapses – indicates that with our n and effect size, we have 100% power to detect a positive FRET signal (effect size 3.8 based on the % Aβ-TMEM97 FRET positive pixels within PSDs 38.4% ± 9.66 and the biological negative control – the % PSD-synaptophysin FRET positive pixels 1.75% ± 0.203). Clinical and neuropathological data were retrospectively obtained from the clinical charts available at the Edinburgh Brain Bank. Neuropathological stages were applied according to international recommendations.[7, 45, 66] Details of the human cases included are found in **Table 1**. Use of human tissue for post-mortem studies has been reviewed and approved by the Edinburgh Brain Bank ethics committee and the ACCORD medical research ethics committee, AMREC (ACCORD is the Academic and Clinical Central Office for Research and Development, a joint office of the University of Edinburgh and NHS Lothian, approval number 15-HV-016). The Edinburgh Brain Bank is a Medical Research Council funded facility with research ethics committee (REC) approval (16/ES/0084).

**Table 1.**
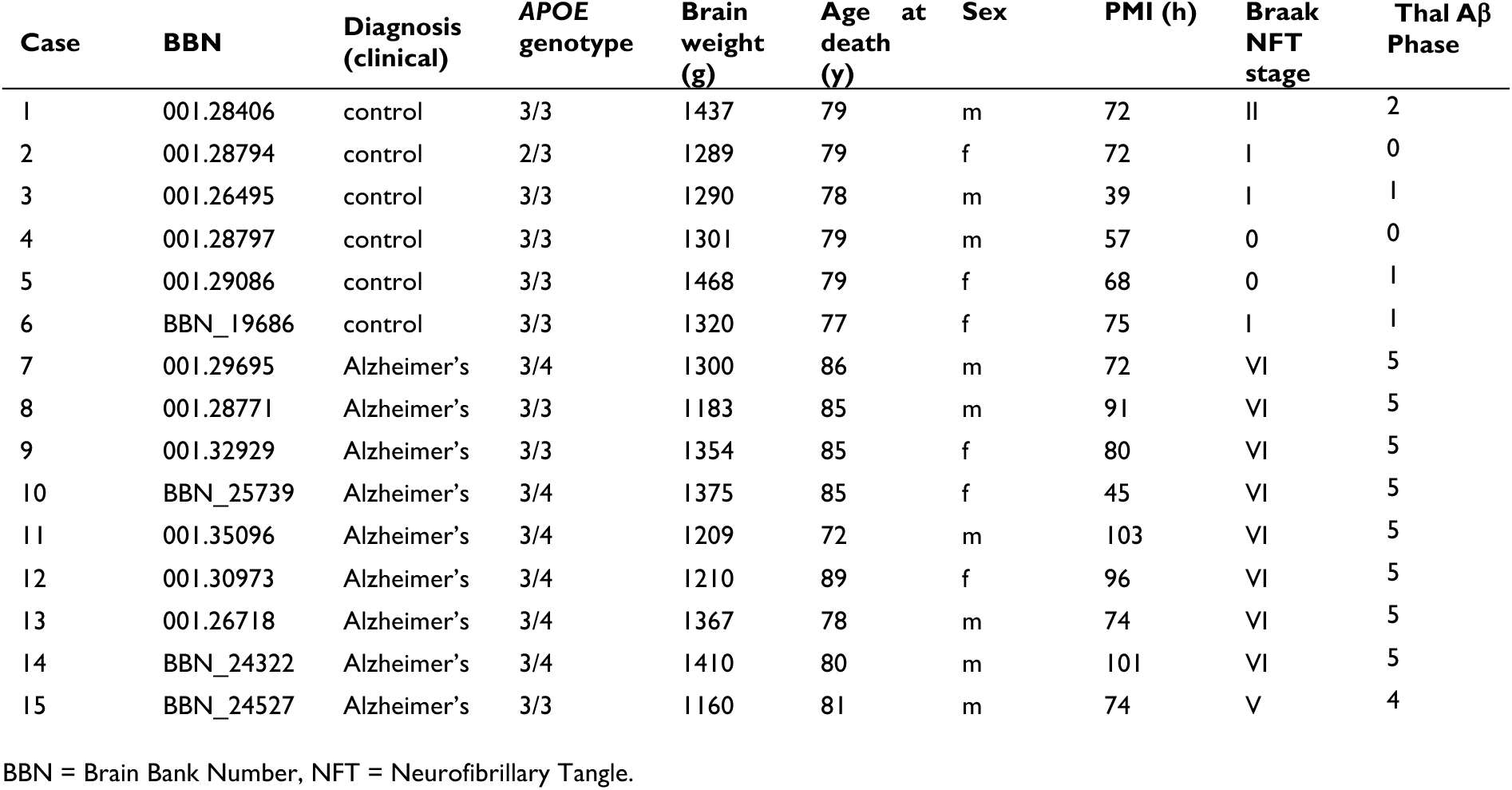
Demographic, clinical, neuropathological and genetic data of human cases.

### Mice

Mice expressing both human tau and the APP/PS1 transgene (APP/PS1+Tau) were generated as previously described.[51] Briefly, two feeder lines were bred to produce experimental genotypes. The feeder lines were line 1: mice heterozygous for an APP/PS1 transgene and a CK-tTA driver transgene and homozygous for knockout of endogenous mouse tau mouse tau; line 2: heterozygous for the Tg21221 human wild type tau transgene driven by CK-tTA and homozygous for knockout of endogenous mouse tau mouse tau.[51] APP/PS1+Tau mice (n=20) and littermate control mice not expressing APP/PS1 nor tau (n=20) were aged to 9 months old before starting CT1812 treatment. Mice of both sexes were randomised into vehicle or control groups. Animal experiments were conducted in compliance with national and institutional guidelines including the Animals [Scientific Procedures Act] 1986 (UK), and the Council Directive 2010/63EU of the European Parliament and the Council of 22 September 2010 on the protection of animals used for scientific purposes, and had full Home Office ethical approval.

Mice were singly housed in a 12 hour dark/light cycle with food and water *ad libitum.* Before dosing started, mice were habituated with double concentration Hartley’s strawberry jelly 4 days during which time all mice learned to eat the entire serving of jelly within 5 minutes. CT1812 fumarate was dissolved in dimethyl sulfoxide (DMSO) and added to cold jelly solution to make up the final volume of 0.6mg/ml concentration before being allowed to set. Each week, a batch of Hartley’s strawberry jelly containing CT1812 or vehicle (plain triple strength jelly) was made. Mice were weighed at the beginning of each week to determine the weight of jelly to be given for that week, and were dosed daily for one month with jelly containing vehicle or CT1812 10 mg/kg/day (experimenters administering jelly were blind to condition). The jelly was delivered in a small petri dish on the floor of the home cage and mice were observed until all jelly was eaten to ensure the full dose was received following which the empty dish was removed.

After 28 days of treatment, mice were sacrificed by terminal anaesthesia. Blood was collected for drug levels then mice were perfused with phosphate buffered saline (0.1M). Brains were removed and the cerebellum snap frozen for testing drug levels. One cerebral hemisphere (selected randomly) was drop fixed in 4% paraformaldehyde. The other hemisphere was dissected and entorhinal cortex processed for array tomography. The rest of the hemisphere was snap frozen for biochemical studies. Estimated percent receptor occupancy was calculated according to the formula (concentration/Ki)/[(concentration/Ki) + 1)], where Ki is determined by radioligand competition binding.[25]

The main study combining array tomography and FRET experiments were performed on APP/PS1+tau mice (n=10) and control littermates of mice (n=8). Details are found in **Table S1**. Standard array tomography imaging (without FRET) was performed on APP/PS1+tau mice (n=9) and control littermates of mice (n=13) to test whether there were any drug effects on synapse density.

### Array tomography

Fresh brain tissue samples from human and mouse cases were collected and processed as previously described.[31] Briefly, small pieces of brain tissue comprising all cortical layers were fixed in 4% paraformaldehyde and 2.5% sucrose in 20mM phosphate-buffered saline pH 7.4 (PBS) for up to 3h. Samples were then dehydrated through ascending concentrations of cold ethanol until embedding into LR White resin (Electron Microscopy Sciences, EMS), which was allowed to polymerize overnight at >50°C. Tissue blocks were then stored at room temperature until used. For each case, two blocks corresponding to BA20/21 for human cases, or one from entorhinal cortex for mouse samples, were cut into 70nm thick sections using an ultramicrotome (Leica) equipped with a Jumbo Histo Diamond Knife (Diatome, Hatfield, PA). Ribbons of at least 20 consecutive sections were collected in gelatine subbed coverslips.

70nm thick ribbons were immuno-labelled as described previously.[31] Briefly, coverslips were first incubated with Tris-glycine solution 5min at room temperature followed by blocking of non-specific antigens with a cold-water fish blocking buffer (Sigma-Aldrich) for 30min. Samples were then incubated for 2h with primary antibodies, washed with Tris-buffered saline (TBS) solution and secondary antibodies applied for 30min. After another TBS washing cycle, coverslips were mounted on microscope slides with Immu-Mount (Fisher Scientific) mounting media. For the detailed information of the primary and secondary antibodies used, please see **Table S2**.

For FRET imaging, images of the same field of view of the consecutive sections were acquired using a Leica TCS8 confocal with 63x 1.4 NA oil objective. Alexa fluor 488, Cy3 or Cy5 were sequentially excited with the 488, 552 or 638 laser lines and imaged in 500 to 550 nm, 570 to 634 nm or 649 to 710 nm spectral windows, respectively. For FRET analysis, the spectral window of the Cy5 (the acceptor, 649 to 710 nm) was also imaged under the excitation of Cy3 (the donor, 552 nm). This setting allowed us to record the transfer of energy from donor molecules to acceptors based on intensity (sensitized emission FRET,[22, 76], **Fig. S1**). Laser and detector settings were maintained through the whole study avoiding major saturation, which is only applied in figures for image visualization purposes.

Standard array tomography imaging (without FRET) was performed on APP/PS1+tau mice (n=9) and control littermates of mice (n=13) to test whether there were any drug effects on synapse density. These images were acquired on a Ziess Axio Imager Z2 epifluorescence microscope with a 63x 1.4 NA oil immersion objective and a CoolSNAP digital camera.

Images from consecutive sections were transformed into stacks using ImageJ.[57, 59] The following steps were performed using an in-house algorithm developed for array tomography image processing and analysis freely available (based on,[12] available at https://github.com/Spires-Jones-Lab, **Fig. S1**). The consecutive images were first aligned using a rigid and affine registration. For the study of the immunoreactivity patterns, semi-automatic local threshold based on mean values was applied specifically for each channel yet common for all the included images. Importantly, only those objects detected in more than one consecutive section (3D objects) were quantified, allowing us to reduce non-specific signals. The number of objects from each channel were quantified and neuropil concentration in mm^3^ of tissue established after removing confounding structures (i.e. blood vessels or cell bodies). In order to investigate the relationship between channels, colocalization was based on a minimum overlap of 10% of the area of the synaptic terminals. Finally, in Alzheimer’s cases, the concentration of objects in each channel or the colocalizing objects were also determined by calculating the Euclidean distance between the centroid of each object and the closest point to the plaque core perimeter. Objects were then binned in 10μm groups.

For FRET analysis, donor-only (Cy3) and acceptor-only (Cy5) samples were imaged in each imaging session in order to calculate the donor emission crosstalk with the acceptor emission (beta parameter) and the direct excitation of the acceptor by the donor excitation laser line (gamma parameter).[71, 76] Aligned stacks of images corresponding to the acceptor emission under donor excitation line (FRET image) were first corrected for the above-mentioned parameters. Each pixel of the FRET image was corrected according to the pixel intensity of either donor-excited donor-emission images or acceptor-excited acceptor-emission images **Fig. S1**). Using the binary masks created before corresponding to postsynaptic terminals, donor and acceptor images, the pixels where the three objects were found overlapping were studied. The percent of pixels where any FRET signal was observed were quantified, allowing us to have a qualitative measure of the occurrence of the FRET effect.

### iPSC to cortical neuron differentiation

iPSC lines derived from peripheral blood mononuclear cells from participants in the Lothian Birth Cohort 1936 (LBC1936) were used for this study as previously described.[33, 67, 72] In this study we used lines EDi030, EDi034, and EDi036. Neuronal differentiation was induced with dual SMAD inhibition (10mM SB431542, [Tocris, 1614] and 1mM dorsomorphin [R&D, 3093/10]) as published previously.[61] After 12 days induction, neuroepithelial cells were passaged mechanically onto 1:100 Matrigel (Corning, 354230) and maintained in N2B27 media (1:1 of DMEM F12 Glutamax [Thermo Fisher Scientific, 10565018] and Neurobasal media [Thermo Fisher Scientific, 12348017], 1X N-2 (Thermo Fisher Scientific, 17502-048], 1X B-27 [Thermo Fisher Scientific, 17504-044], 1mM L-Glutamine [Thermo Fisher Scientific, 25030-024], 5mg/mL insulin [Merck, I9278-5ML], 100mM 2-mercaptoethanol [Thermo Fisher Scientific, 31350010] 100mM non-essential amino acids [Thermo Fisher Scientific, 11140-050]), and 1X anti-biotic/anti-mycotic [Thermo Fisher Scientific, 15240062]). Neural precursor cells were passaged with accutase (Thermo Fisher Scientific, A11105-01) at day 20, and day 25. A final passage was performed at day 30, with cells plated onto poly-L-ornithine (Merck, P4957) treated glass cover slips coated with 1:100 Matrigel, 10mg/mL laminin (Merck, L2020-1MG), and 10mg/mL fibronectin (Merck, F2006). Between days 35-49 maturing neurons N2B27 was supplemented with 10mM forskolin (Tocris, 1099). From day 50 on N2B27 was supplemented with 5ng/mL BDNF (R&D Systems, 248-BD) and 5ng/mL GDNF (R&D Systems, 212-GD).

Generation of brain homogenate from Alzheimer’s patients to challenge iPSC neurons was conducted according to a published protocol (Hong et al., 2018) with modifications. Human brain tissue was homogenised with a Dounce homogeniser and placed in a low protein binding 15mL tube (Thermo Fisher Scientific, 30122216) containing 10mL 1X artificial CSF (pH 7.4) supplemented with 1x cOmplete mini EDTA-free protease inhibitor cocktail tablet (Roche, 11836170001) per 10mL, per 2g of tissue. The solution was placed on a roller for 30 mins to extract soluble proteins, then centrifuged at 2000 RCF for 10 mins to remove large, insoluble debris. The supernatant was transferred to ultracentrifuge tubes (Beckman, 355647) and then centrifuged at 200,000 RCF for 110 mins. The resulting supernatant, a homogenate fraction containing soluble Aβ forms, was then transferred to a Slide-A-Lyser G2, 2K MWCO 15mL dialysis cassette (Thermo Fisher Scientific, 87719) and dialysis was conducted in 1X aCSF with magnetic stirring for three days at 4°C to remove salts from the homogenate. During this time, the 1X aCSF was exchanged every 24 hours. Dialysed brain homogenate was divided into two equal portions in low protein binding 15mL tubes. Protein A Agarose (PrA) beads (Thermo 20334) were washed three times in 1X aCSF. 30uL of Washed beads were then added per 1mL of homogenate. To create Ab-treatment samples, Aβ was immunodepleted by adding 20µL 4G8 antibody (Biolegend, 800711) per 1mL of homogenate and 20µL 6E10 antibody (Biolegend, 803001) per 1mL of homogenate. To create Ab+ treatment samples, homogenate was ‘mock-immunodepleted’ with isotype control antibodies to non-human antigens by adding 20µL of GFP (DSHB, DSHB-GFP-12A6) and GFP (DSHB, N86/38) antibody per 1ml of homogenate. Concentration of Aβ_42_ was determined by ELISA (WAKO 4987481457102). Homogenate was then incubated for 24 hours on a rocker, during which time the Aβ antibody complexes bind to the PrA beads in the immunodepleted portion. After 24 hour incubation, homogenate was centrifuged at 2500 RCF for 5 mins to remove the beads, and the supernatant was collected. The process of adding beads and antibody/serum to homogenate was repeated twice more. After the third centrifugation step, PrA beads alone were added to both Aβ+ and Aβ-homogenate, incubated for two hours on a rocker, and then centrifuged at 2500 RCF for 5 mins to clear any remaining antibody. Finally, homogenate from each portion was aliquoted at 0.5mL into 1.5mL low protein binding Eppendorf tubes (Thermo Fisher Scientific, 0030108116) and stored at –80°C. Concentration of Ab1-42 in Aβ+ and Aβ-homogenate was quantified by sandwich ELISA (WAKO), according to manufacturer instructions using a ClarioSTAR spectrophotometer (BMG Labtech).

To determine whether Aβ treatments induce cell death, Click-iT™ Plus TUNEL Assay Kits for In Situ Apoptosis Detection (Thermo Fisher Scientific, C10617) was used to detect apoptotic cells according to the manufacturer’s protocols. Samples were fixed with 4% formalin (Polysciences, 04018-1), permeabilized with 0.3% Triton X-100 in 1X PBS for 20 minutes at room temperature, incubated with TUNEL reaction buffer for 10 minutes at 37°C, incubated with TUNEL reaction mixture for 1 hour at 37°C, blocked with 3% bovine serum albumin, and incubated with TUNEL reaction cocktail for 30 minutes at 37°C. Immunocytochemisty was then performed for co-staining. All incubations were conducted in the dark.

Neurons from 3 iPSC donors were grown to approximately day 200 post-induction in 24 well plates. Cells were treated with media, Aβ+ homogenate, or Aβ-homogenate for 24 hours followed by addition of CT1812 (10 mM) or DMSO (Merck, D2438-50ML) vehicle treatment for a further 24 hours. RNA was harvested from four pooled wells per treatment condition using trizol-chloroform extraction. Remaining coverslips were fixed for immunocytochemistry (ICC) as below. Each experiment was repeated with 3 different differentiations of each of the three lines.

Cells for ICC were fixed with 4% formalin (Polysciences, cat.04018-1) for 15 mins, Washed thrice in 1X phosphate buffered saline (PBS). Fixed cells were permeabilized and blocked with in 1X PBS with 0.3% Triton-X and 3% bovine serum albumin (permiabilising block solution) for 30 minutes. Coverslips were incubated overnight at 4°C with primary antibodies TMEM97, homer1, MAP2, GFAP, and Tuj1. Cells were washed with 1X PBS, and incubated in secondary antibodies diluted 1:500 in permeabilising block solution for 1 hour in the dark. For the detailed information of the primary and secondary antibodies used, please see **Table S2**. Cells were incubated with 1:10,000 DAPI in 1X PBS with 0.3% Triton-X for 10 minutes, and washed 2x in 1X PBS. Coverslips were mounted on slides (VWR, 631-0847) with mounting media (Merck, cat.345789-20ML) and imaged on a Leica TCS confocal microscope with an oil immersion 63x objective.

RNA sequencing was performed on total RNA samples using TruSeq stranded mRNA-seq library preparation along with next-generation sequencing on NovaSeq6000 platform; sequencing was carried out by Edinburgh Genomics (Edinburgh, UK). Samples were sequenced to a depth of approximately 100 million 50-base pair, paired-end reads. The reads were mapped to the primary assembly of the human (hg38) reference genome contained in Ensembl release 106, using the STAR RNA-seq aligner, version 2.7.9a.[17] Tables of per-gene read counts were generated from the mapped reads with featureCounts, version 2.0.2.[39] Differential gene expression was performed in R using DESeq2, version 1.30.1.[40] Gene ontology analyses were run on the Gene Ontology online resource using their Panther online search tool for Biological Processes (http://geneontology.org/). MetaCore+MetaDrug version 22.3 build 71000 was used to perform pathway analysis on Abeta vs. Vehicle, and Abeta + Drug vs. Abeta + Vehicle conditions (un-adjusted p-value<0.05). STRING (Version 11.5) pathway analysis of Abeta + Drug vs. Abeta + Vehicle conditions (un-adjusted p-value<0.05).[77]

### Statistical analysis

Brain weight, age at death, and PMI differences between groups were analysed with t-test or Wilcoxon test depending on the Shapiro-Wilk Normality test results. Sex, *APOE* genotype, Braak stage, and Thal phase were analysed with Fisher-exact test. The comparison between groups in all the other studied variables was analysed using linear mixed effects models including case or cell line as a random effect to account for multiple measures. Sex, age, APOE4 status, Braak stage, Thal phase and PMI were included as a fixed effects in the initial statistical models for human post-mortem data followed by sequential removal of fixed effects to find the model that best fit the data for our primary question of whether Aβ and TMEM97 are found within the same synapses. The linear mixed effects model including only diagnosis as a fixed effect and sample nested in case as a random effect was the best fit for the data as assessed with the Akaike Information Criteria (AIC).[9] Despite the model including sex being a slightly poorer fit, we appreciate that inclusion of sex as a biological variable is best practice in dementia research[42] so sex remained in the model. Including an interaction between sex and diagnosis was a better fit than without the interaction, thus our final statistical model applied in the study was diagnosis*sex+(1|case/sample). For analysis of the effect of plaque promximity, distance from plaque was included as a fixed effect in the model. ANOVAs and Tukey corrected post-hoc were run on the linear mixed effects models to determine differences between groups. All the analyses were performed with R v4.1.2 [53] and the scripts and full statistical results can be found at *add upon acceptance for publication*.

### Study design

The immunostaining, image acquisition, image processing and analyses were performed blinded to the clinicopathological diagnosis. Mice were randomly assigned to treatment groups and treated by blinded experimenters. Bias was also minimized by setting the same parameters for image acquisition and image analysis for all the included cases.

### Data availability

Protocols, image analysis scripts and R scripts for statistical analysis are available at *link to be added upon acceptance for publication*. Raw images available from the corresponding author upon reasonable request.

## Results

### Characteristics of human cases

We used human post-mortem brain samples from inferior temporal gyrus (BA20/21) to investigate proximity of Aβ and synaptic proteins. Details of human cases included are shown in **Table 1**. Our Alzheimer’s and control cohorts are age and sex matched (*p*>0.05, Wilcoxon and Fisher-exact test, respectively). *APOE* e4 carriers were more common in Alzheimer’s than in the control group (Fisher-exact test, *p*=0.028). Post-mortem interval (PMI) was slightly longer in Alzheimer’s group (Welch’s t-test, *p*=0.04975). Both Braak stage and Thal phase were higher in AD than control group due to our inclusion criteria for the groups (p<0.01 Fisher’s exact tests.

### TMEM97 levels are increased in Alzheimer’s

We used array tomography to examine synaptic localisation of Aβ, TMEM97, and other potential binding partners. Image stacks were acquired in areas containing amyloid plaques where we previously demonstrated the highest amount of synaptic Aβ. The overall density of TMEM97 positive objects was assessed in the temporal cortex revealing an immunoreactivity pattern of a membrane protein, with widespread presence in grey matter in both Alzheimer’s and control cases (**Fig. 1A**). The density of TMEM97 objects was significantly higher in Alzheimer’s than control cases (fold increase: 1.52; effect of disease F[1,10.73]=5.53, p=0.039, no significant sex differences). This increase was not related to the proximity of an Aβ plaque (**Fig. 1B**). As previously described [34], postsynaptic terminal density was reduced in the vicinity of Aβ plaque cores of Alzheimer’s cases (effect of plaque distance F[4,Inf]=5.69, p=0.0001), although no overall loss was observed when compared with control cases (**Fig. 1C**). As expected, Aβ was more common in Alzheimer’s cases and the density of objects was elevated close to the core of Aβ plaques (effect of plaque distance F[4,Inf]=34.45, p<2E-16; **Fig. 1D**).

**Figure 1.**
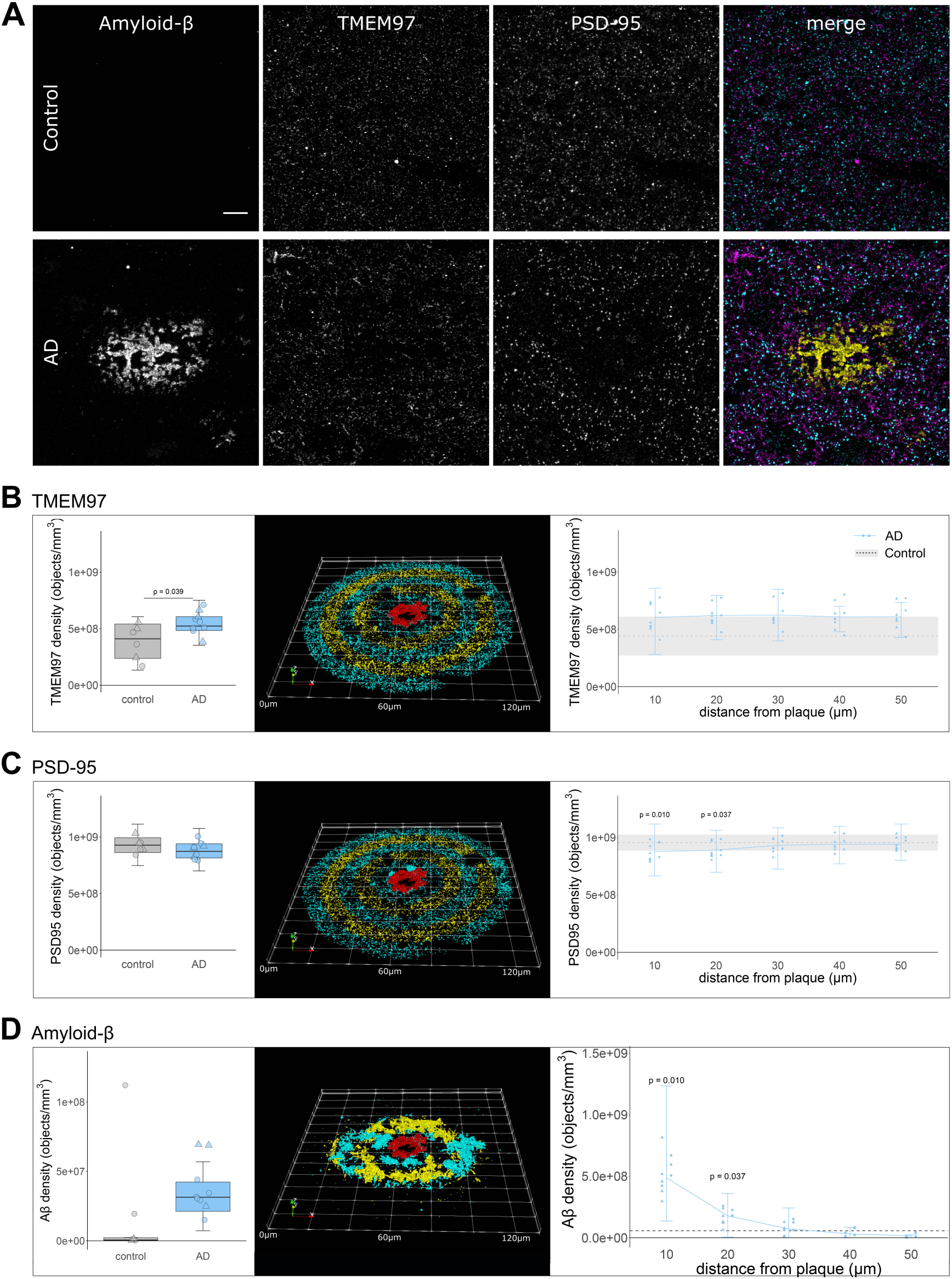
Immunoreactivity pattern and density of TMEM97, Aβ and PSD95. (**A**) representative maximum intensity projection images of 10 consecutive 70nm-thick sections from a control and an Alzheimer’s case. Immunoreactivity against Aβ (yellow), TMEM97 (magenta) and PSD95 (cyan) is shown. Overall density (left) or the density in relation to Aβ plaque cores (right) of TMEM97 (**B**), PSD95 (**C**) and Aβ (**D**) is plotted. The 3D reconstructions were made from 19 consecutive sections of a representative Alzheimer’s case. The Aβ core is shown in red and the objects distributed every 10um bins are coloured. Scale bar represents 10µm. Boxplots show quartiles and medians calculated from all image stacks in the study. Data points show case means (females, triangles; males, circles). P values on left panels show significant effect of disease (ANOVA after linear mixed effects model). On right panels, p values show Tukey corrected post-hoc significant differences between 50 μm and the indicated plaque distance in the AD data. In D, note the scales are different in the two plots as there is an order of magnitude more Aβ near plaques than when averaged across all images.

### TMEM97 is found in a higher proportion of synapses in Alzheimer’s

We recently described an increase of TMEM97 protein levels in biochemically isolated synaptic fractions from Alzheimer’s brain compared to controls.[24] In the present study, we were able to visualize the synaptic localization of TMEM97. The analysis of the 1,112,420 single synaptic terminals revealed an increased proportion of synapses with TMEM97 in Alzheimer’s when compared to healthy controls (fold increase: 1.77; effect of disease F[1,10.27]=12.59, p=0.005, **Fig. 2B**). There were no sex effects in the proportion of synapses containing TMEM97. In line with the hypothesis that TMEM97 is a binding partner of Aβ, we found that Aβ was present in postsynaptic terminals (effect of disease F[1,21.55]=9.14, p=0.006, **Fig. 2C**) and, importantly, that Aβ was found overlapping TMEM97 immunoreactivity at the same synapses effect of disease F[1,21.4]=13.74, p=0.001, **Fig. 2D**). Interestingly, there was a significant interaction between disease and sex in the proportion of synapses with Aβ (disease* sex interaction F[1,21.55]=7.2, p=0.013) and those with Aβ and TMEM97 (disease* sex interaction F[1,21.4]=9.0, p=0.007) with female subjects having more synaptic Aβ.

**Figure 2.**
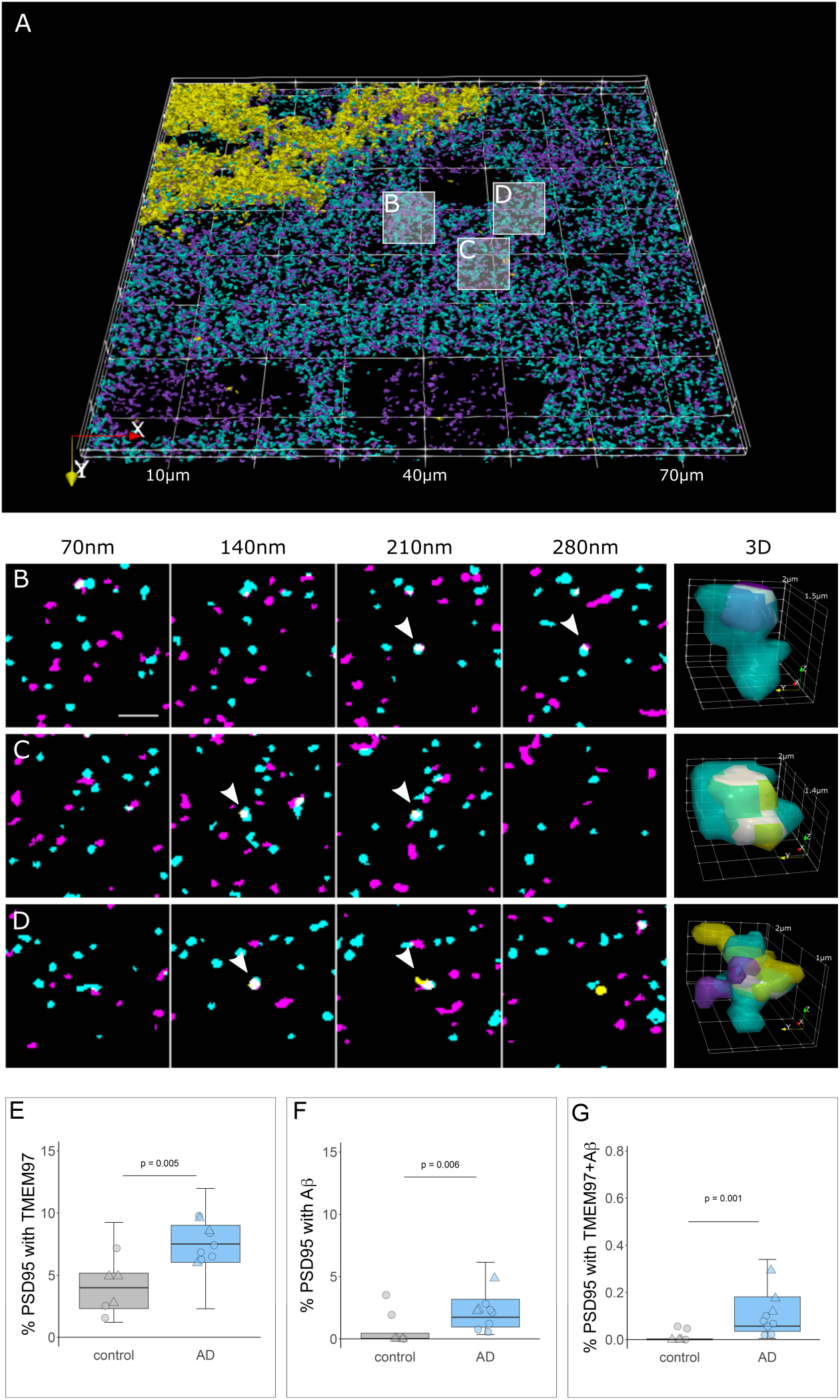
TMEM97 is found at higher levels in Alzheimer’s synaptic terminals compared to healthy controls. 3D reconstructions were made from 20 consecutive 70nm-thick sections from a representative Alzheimer’s case stained for Aβ (yellow), TMEM97 (magenta) and PSD95 (cyan). In the top 3D reconstruction **(A**), three white boxes label the magnified regions that highlight: a PSD95 terminal with TMEM97 (**B**), a postsynaptic terminal with Aβ (**C**) and a PSD95 synaptic terminal with both Aβ and TMEM97 (**D**). In magnified images **B-D**, four consecutive sections from the image stack are shown (each 70nm apart). White arrowheads indicate synaptic localization and a 3D reconstruction (right panel) of the pointed synapse where colocalization is observed (white). Box plots show the percent of postsynaptic terminals that contained TMEM97 (E), Aβ (F), or both (G), in Alzheimer’s and control cases. Boxplots show quartiles and medians calculated from each image stack. Each data points refers to the means of a single human tissue donor (females, triangles; males, circles). P values show ANOVA after linear mixed effects models. Scale bar: 2µm.

### Synaptic TMEM97 and Aβ are close enough to generate a FRET signal

After confirming the presence of TMEM97 together with Aβ in synapses of Alzheimer’s cases, and given the ability of Sigma-2 receptors to displace Aβ from neuronal synapses,[25] we assessed the proximity of the immunoreactivity of TMEM97 with that of Aβ by FRET. In this single pixel analysis, those areas where the donor (Cy3 labelling Aβ) and the acceptor (Cy5 labelling TMEM97) were found overlapping within a PSD95 positive object were quantified in the corrected donor excitation-acceptor emission image (**Supplementary** Fig. 1 and methods for further details). This quantification allowed us to detect FRET only when both donor and acceptor were present and within approximately 10nm of each other. To determine limitations of the technique, we measured the residual FRET signal in stacks where only the donor or the acceptor was labelled as negative controls, and the maximum FRET signal was determined where the donor and acceptor labelled the same target as the positive control (**Fig. 3**, green shading on the plot shows the experimental window of detecting FRET signal between negative and positive control levels).

**Figure 3.**
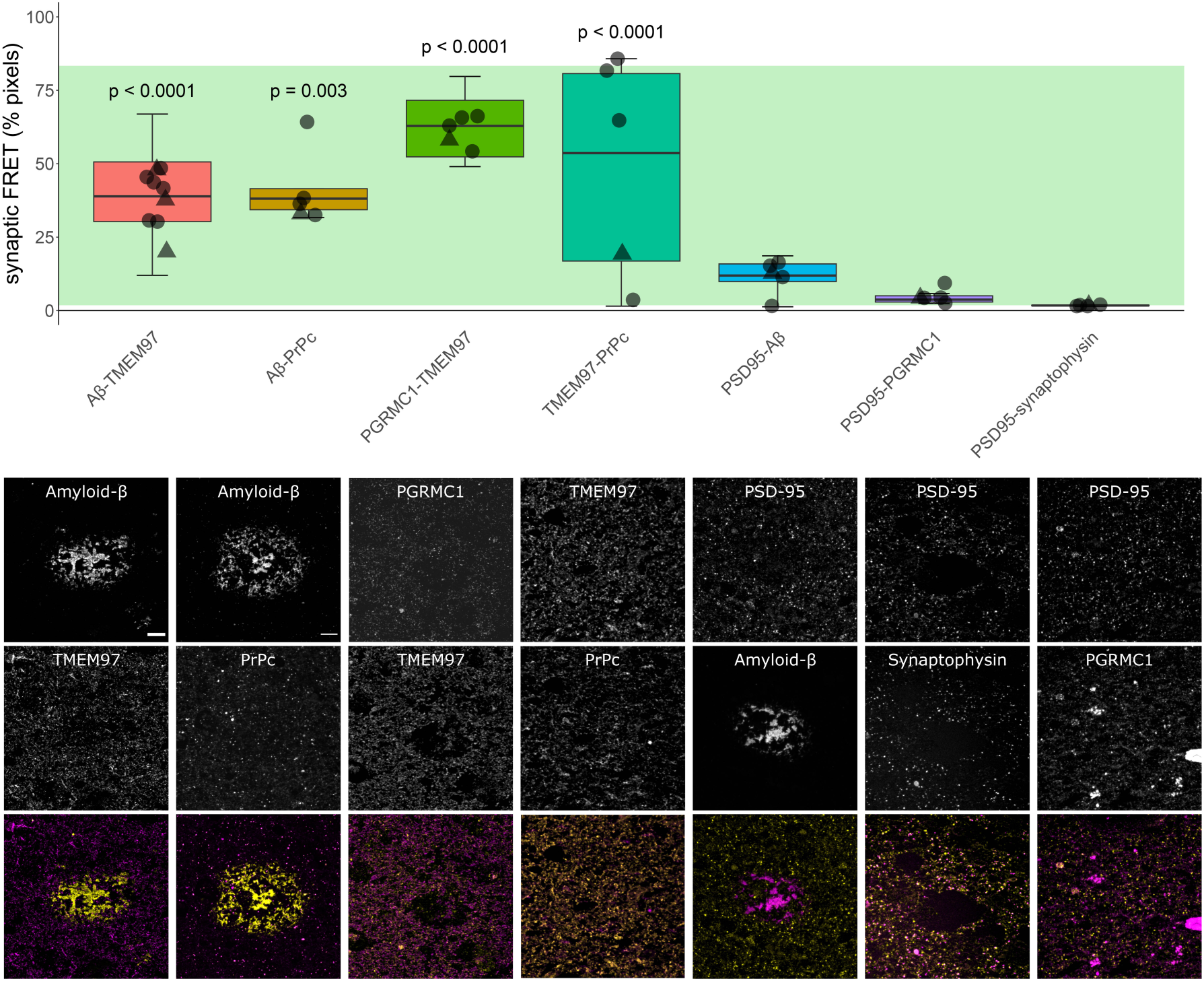
Aβ and TMEM97 are close enough at the Alzheimer’s synapses to generate a FRET effect. Pixels where pairs of interest were colocalized within a PSD95 puncta were analyzed to determine whether they generate a FRET signal. The percent of synaptic pixels where FRET signal was detected by each protein pair are plotted (top panel). The green bar in the boxplot shows the window of detecting FRET signal defined by the positive control signal where an acceptor fluorophore was applied to the same protein as the donor fluorophore using a tertiary antibody (top) and negative controls where no acceptor fluorophore was present (bottom). Boxplots show quartiles and medians calculated from each image stack. Data points show case means (females, triangles; males, circles). P-values show post-hoc Tukey corrected differences between the pair indicated and the biological negative control of PSD-synaptophysin FRET. Images below the graph show maximum intensity projections of 10 consecutive 70nm-thick sections of the protein pairs tested for FRET effect. Each targeted protein is shown individually in grayscale and in the merged images at the bottom, the donor is shown in yellow and the acceptor in magenta. Scale bar: 10µm. Abbreviations: PrP_c_, cellular prion protein; Syph, synaptophysin.

In Alzheimer’s cases, we found that on average, in 38.4±9.66% of synaptic pixels where donor and acceptor were present, Aβ and TMEM97 were close enough to generate a FRET signal (**Fig. 3**). We also observed FRET between Aβ and PrP_c_ – which has also been observed to bind Aβ in model systems[62] – and between TMEM97 and PGRMC1 which are known to be binding partners in vitro and in human cases [55], and TMEM97 and PrP_c_ (**Fig. 3**). Lastly and confirming cellular localization of Aβ to synapses, we saw positive FRET signal between Aβ and PSD95 (**Fig. 3**). This is in support of findings by other groups in which Aβ has been described to interact in some synapses, including in rat neuronal cultures [28] and our previous study using a similar FRET approach in APP/PS1 mice.[35, 50, 52]

To confirm that this effect was not occurring in all areas where donor and acceptor are present in the same pixel, we used a biological negative control looking for FRET between PSD95 and synaptophysin which are close but not interacting as they are separated by the synaptic cleft. As expected, there was not a significant FRET signal between these pre and postsynaptic proteins. There was also no FRET signal between PGRMC1 and PSD95 (**Fig. 3**). In summary, our FRET experiments confirm close proximity of Aβ and TMEM97, Aβ and PrP_c_, TMEM97 and PrP_c_, and TMEM97 and PGRMC1, and Aβ and PSD95, but not PSD95 and synaptophysin, and PSD95 and PGRMC1, which are robust as both technical and biological negative controls do not show FRET signal.

### TMEM97 modulator reduces synaptic TMEM97-Aβ FRET signal in a mouse model of Alzheimer’s

Results from human brain observations suggested that TMEM97 may be a binding partner of Aβ. To determine whether this synaptic binding is reversible in vivo, we used the Sigma-2/TMEM97 receptor complex allosteric modulator CT1812 – currently in clinical trials for Alzheimer’s[11, 20] – in a recently described Alzheimer’s mouse model.[11, 20, 51] APP/PS1+Tau mice (APP-PS1+/–; MAPT –/–; CKTTA +; Tg21221) and littermate controls were treated with either vehicle (n=10 APP/PS1+Tau, n=10 control) or CT1812 (10mg/kg/day given orally, n=10 APP/PS1+Tau, n=10 control), which selectively binds to the Sigma-2 (TMEM97) receptor complex.

We first estimated the percent receptor occupancy of CT1812 in brain, which was calculated based on the measured brain concentration of the drug (see methods and [25]). We observed a statistically significant sex difference in the percent of estimated receptor occupied by the drug. Male APP/PS1+Tau mice had an average of 85.13±6.4% estimated receptor occupancy, while females had an average of 69.69±11% (β=15.44; p<0.001, **Fig. 4B**). The increase in drug estimated receptor occupancy in male mice was observed in both genotypes (β=44.63; *p=*0.006, **Supplementary** Fig. 2C). This difference could not be explained by any experimental procedures as all animals were given the same dose of compound from the same stock. Body weight, which could affect drug uptake, did not differ between treatment groups (F[1,36]=0.74, *p=*0.40), but body weight was different between male and female mice with males being heavier (F[1,36]=62.33, p<0.0001). There was no treatment:sex interaction in analysis of body weight nor was there a significant difference in the numbers of male vs female mice in the different treatment groups (Fisher test *p=*1, 95% CI = 0.30, 5.04). The treatment of non-transgenic control mice did not affect the density of Aβ, Tau or PSD95 (**Supplementary** Fig. 2A-B), confirming that treatment with this compound was not synaptotoxic. CT1812 treatment in transgenic mice did not change plaque burdens but did reduce astrogliosis in hippocampus (Supplementary Fig. 3).

**Figure 4.**
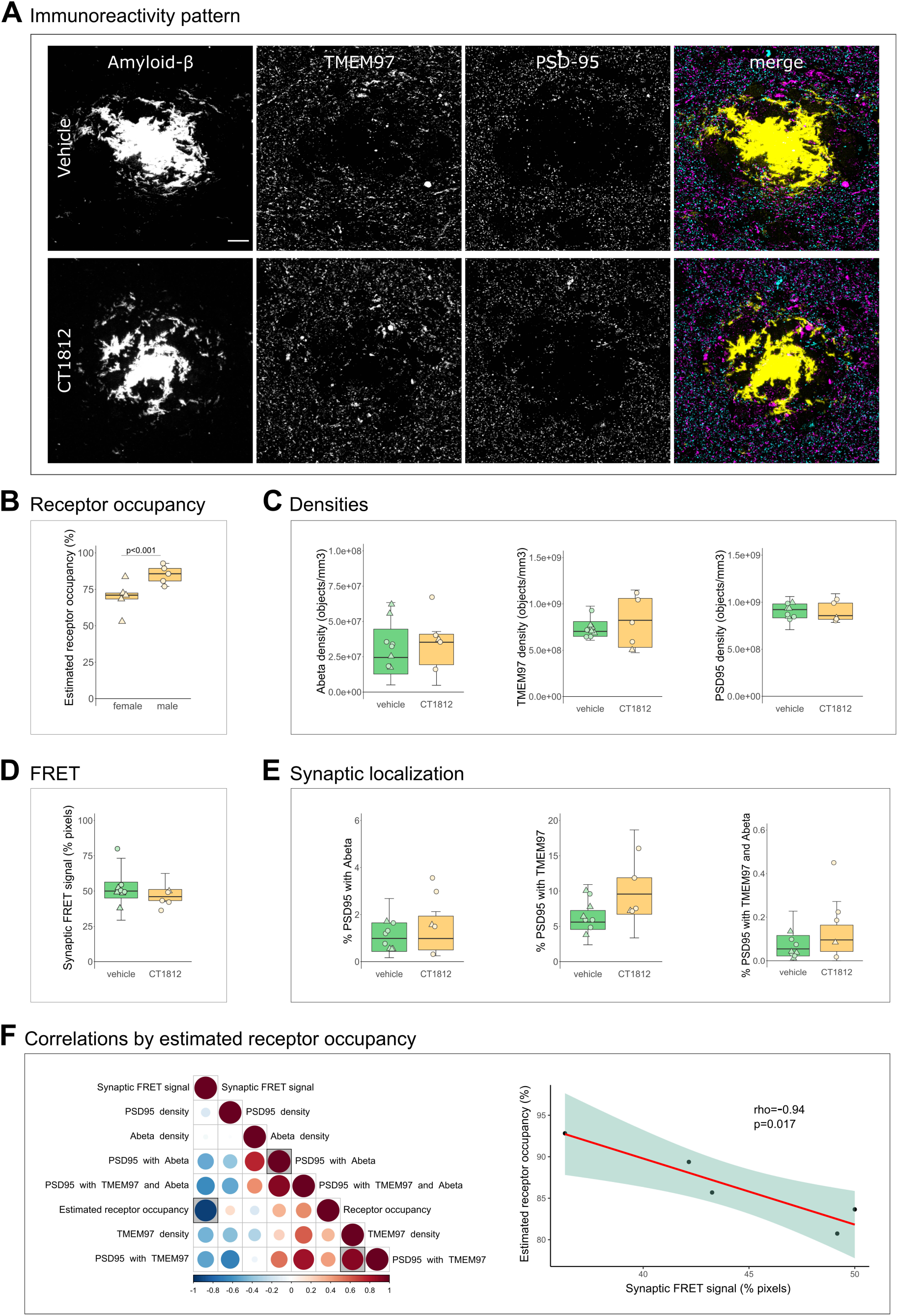
Effect of TMEM97 modulator on synaptic Aβ and TMEM97 in the APP/PS1+Tau mice model. Representative images of immunoreactivity patterns found in vehicle or CT1812 treated mice are shown in **A**. Images show maximum intensity projections of 16 consecutive 70nm-thick sections of cases stained for Aβ (yellow), TMEM97 (magenta) and PSD95 (cyan). **B**, the estimated percent of receptor occupancy by the drug in the CT1812 treated group. **D**, the percent of synaptic pixels that contain both Aβ and TMEM97, and FRET signal. **C,** quantification of overall densities of the three studied proteins. **D**, the postsynaptic terminals localisation of Aβ, TMEM97, or both. **F**, Correlations were estimated between measured parameters and a correlation matrix of the assessed variables is shown (left panel) in which the colour and size reflect the rho (scale below the plot) and the statistically significant correlations are highlighted with a shaded square. The correlation between percent estimated receptor occupancy and percent of synaptic FRET signal (right panel) displaying the regression line (red), the 95% confidence interval (green) and the Spearman correlation results (rho, p value). Scale bar: 10µm. Boxplots show quartiles and medians calculated from each image stack. Data points refer to case means (females, triangles; males, circles). Analysis with linear mixed effects models including treatment group and sex interaction.

Since it has been reported that drug concentrations above 80% estimated receptor occupancy are effective and concentrations lower than 60% were ineffective,[25, 26] the effect on synaptic TMEM97 and Aβ localization was studied on the mice that reached an 80% estimated receptor occupancy threshold (n=5 APP/PS1+Tau mice, **Table S1**). CT1812 did not statistically significantly influence the overall densities of Aβ, TMEM97 or PSD95 nor the synaptic localization of Aβ and/or TMEM97 (**Fig. 4A, C, E**). When we modelled the effect of treatment and sex on the synaptic FRET signal between Aβ and TMEM97, we did not observe a significant difference between groups (vehicle mean 52.8±12%; treated mean 44.2±5.61%, **Fig. 4D**). However, the increase of estimated receptor occupancy by the drug significantly correlated with a decrease of synaptic FRET signal between Aβ and TMEM97 (*rho*=-0.94, *p=*0.017, **Fig. 4F**).

Taken together, we found that in the CT1812 treated APP/PS1+Tau mice with estimated receptor occupancy in the therapeutic range, there was a decreased synaptic FRET signal between Aβ and TMEM97, indicating increased distance between the two proteins.

### TMEM97 modulator ameliorates Aβ-induced phenotypes in human iPSC neurons

To investigate whether disrupting TMEM97-Aβ interactions protects living human neurons, human iPSC derived neurons were challenged with the soluble fraction of Alzheimer’s brain homogenate or identical homogenate immunodepleted for Aβ and treated with CT1812 or vehicle to investigate whether disrupting TMEM97-Aβ interactions protects living human neurons. Brain homogenates were added to neuronal media at final concentrations of 90 pM in the mock-immunodepleted condition and approximately 8 pM in the immunodepleted condition (detection was at the lower limit of ELISA sensitivity). Immunocytochemistry confirms that TMEM97 is present in homer positive post synaptic puncta along dendrites and that when treated with Aβ-containing brain homogenate, Aβ also accumulates in synapses (**Fig. 5A**). Both cell counts and TUNEL assay showed that the brain homogenate treatments were not cytotoxic compared to media (**Fig. 5B, C**), which is important as we wish to model synaptic damage, not neuron death, since oligomers of Aβ at physiological concentrations in human brain are thought to cause synaptic damage and not to induce neuron death directly. Cytotoxicity was generally low (<10%) in all conditions.

**Figure 5.**
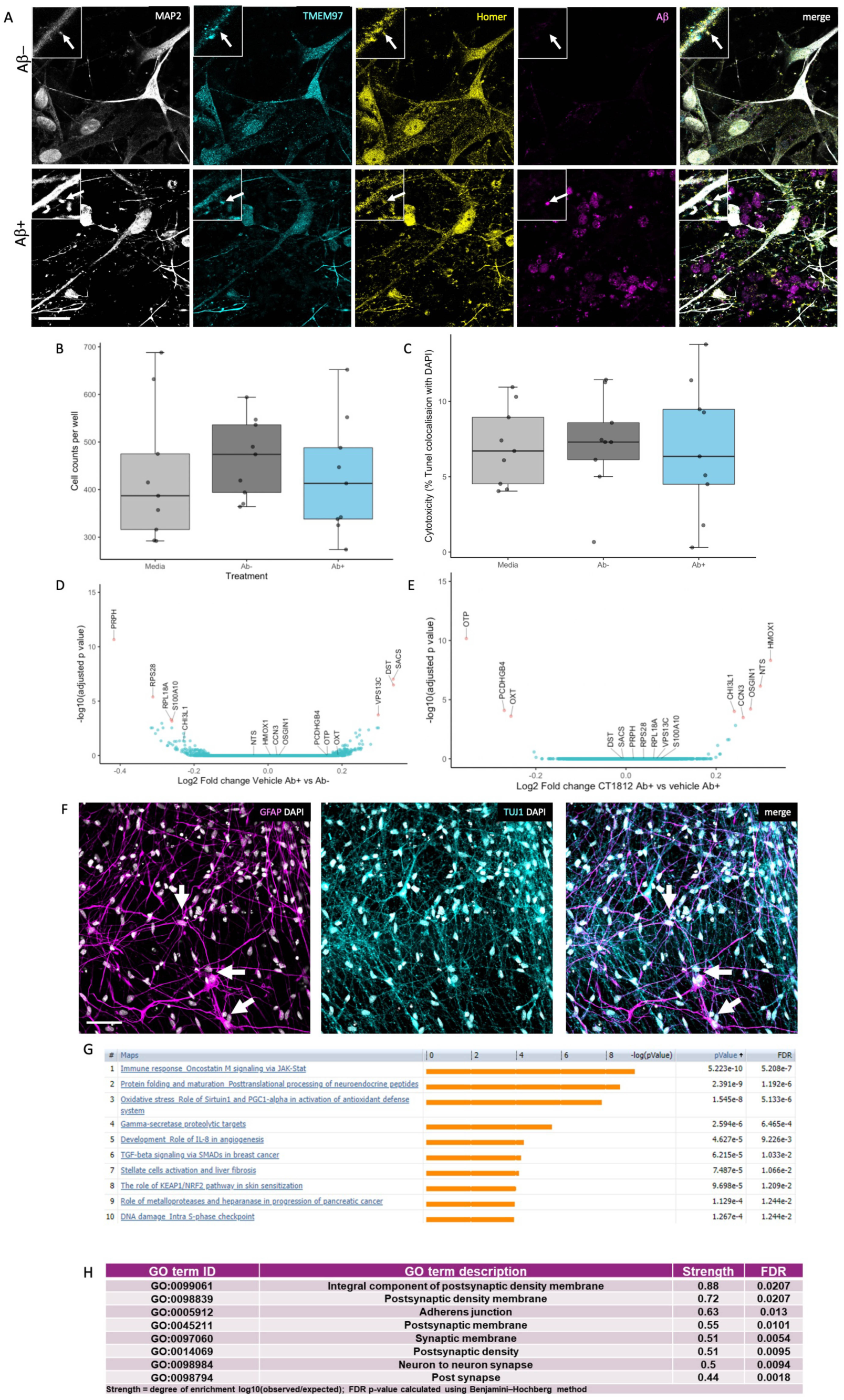
Challenge of human iPSC neurons with Alzheimer’s brain homogenate. (**A**) Immunocytochemistry for dendrites (MAP2, grey), TMEM97 (cyan), postsynapses (homer, yellow), and Aβ (magenta) reveals that Aβ accumulates in TMEM97-containing post synapses in iPSC-derived human neurons challenged with Alzheimer’s brain homogenate when the homogenate was mock immunodepleted (Ab+), but not when it was immunodepleted for Aβ (Ab-). DAPI positive cell counts (**B**) and a TUNEL cytotoxicity assay (**C**) show that brain homogenate treatments do not induce cell death. RNA sequencing reveals 7 significantly differentially expressed genes between Ab+ and Ab–homogenate treatment (**D,** differentially expressed genes with adjusted p value <0.05 shown in pink in volcano plot). When comparing Aβ challenged cultures treated with CT1812 and vehicle, 8 genes are differentially expressed (volcano plot in **E** differentially expressed genes with adjusted p value <0.05 shown in pink). Staining for astrocytes (GFAP, magenta), neurons (TUJ1, cyan), and nuclei (DAPI, white) reveals that a small proportion of cells in our cultures are astrocytes (arrows) which extend many processes (**F**). Pathway analysis using MetaCore (unadjusted p-value<0.05) shows enrichment of immune/inflammatory pathways with Aβ treatment compared to immunodepleted Aβ treatment (**G**). Using STRING protein interaction analysis of AΒ + Drug vs. AΒ + Vehicle conditions (unadjusted p-value<0.05), the top GO Components (sorted by strength) were involved in synaptic biology (top 8 shown, **H**). Scale bar in panel **A** 20 μm, insets 10×10 μm, Scale bar in panel **F** 50 μm.

RNA sequencing was used to determine gene expression changes after exposing cells to human Alzheimer’s brain homogenate containing soluble Aβ and CT1812 or vehicle. Treatment with Alzheimer’s brain homogenate containing soluble Aβ induced over 4,000 differentially expressed genes compared to media treatment. The difference between Aβ– and Aβ+ homogenate was much smaller with only 7 differentially expressed genes (adjusted p-value <0.05), 3 upregulated and 4 downregulated (**Fig. 5D**). Treatment with CT1812 did not significantly change expression of these 7 genes compared to vehicle after Aβ + homogenate. Although not statistically significant, the fold changes of these 7 genes largely changed in the opposite direction with treatment with CT1812 (**Table S3**). Three of the genes upregulated by Aβ + homogenate challenge are expressed in neurons and implicated in neurodegenerative diseases (DYS – dystonin, SACS – sacsin, and VPS13C – vacuolar protein sorting 13). 8 genes were differentially expressed with CT1812 treatment after Aβ + homogenate challenge (**Fig. 5E, Supplementary Table 3**). Several of the transcripts regulated by CT1812 are expressed in astrocytes, which are observed in small numbers in our cultures (averaging 16% GFAP positive cells in the 200 day old cultures used, **Fig. 5F**). One of the transcripts significantly changed by CT1812 treatment, CHI3L1, which encodes YKL-40 protein, is expressed in astrocytes and is an interesting biomarker of inflammation in Alzheimer’s disease.[48]

Although only a handful of transcripts reached a statistically significant change after correcting for multiple testing, pathway analysis using a less strict criteria of *p*<0.05 unajdusted *p*-values to look for patterns of expression changes using MetaCore, revealed enrichment of immune/inflammatory pathways with Aβ treatment compared to immunodepleted Aβ treatment. These included the JAK/STAT pathway known to be involved in astrocyte differentiation and function, indicated by Leukemia Inhibitory Factor (LIF) and Leukemia Inhibitory Factor Receptor (LIFR) enrichment, a known pro-inflammatory cytokine involved in the differentiation of neuronal precursor cells into astrocytes (**Fig. 5F**). STRING (Version 11.5) protein interaction analysis of Aβ + Drug vs. Aβ + Vehicle conditions (unadjusted *p*–value<0.05) confirmed the findings of TMEM97 localization to synapses. The top 8 GO Components (sorted by strength) were involved in synaptic biology (**Fig. 5G**). Further, “Synapse” was the top UniProt Keyword of the Aβ + Drug vs. Aβ + Vehicle (unadjusted p<0.05), with a strength of 0.54 and a FDR of 0.00019. Further, biological pathways changed by CT1812 vs vehicle treatment after challenge with Aβ homogenate include several processes important for synaptic function (**Supplementary Table 4**). Overall, differential expression data and pathway analysis support a localization of – Sigma-2 receptor at the synapse, and suggest a role for CT1812 in modulating inflammatory pathways, perhaps indirectly by glia that may detect changes in synapse health or activity.

## Discussion

In the present study, we visualized TMEM97 within individual postsynapses in human brain. In Alzheimer’s brain, TMEM97 levels were increased and in synapses, and TMEM97 was found in close enough proximity to Aβ to be binding. Further we confirm in mouse and human iPSC models that Aβ is in close proximity to TMEM97 in synapses and that CT1812 treatment to disrupt this interaction is beneficial.

TMEM97 (Sigma-2) has been previously linked to Alzheimer’s, through *in vitro* and *in vivo* pharmacological modulation studies and with regard to a change in expression levels on synapses in Alzheimer’s patients. More specifically, in cellular models, treatment with modulators or knocking out sigma-2 receptor (S2R) constituents TMEM97 or the co-receptor PGRMC1 reduces the internalization of Aβ.[26, 54] In an animal model with Alzheimer’s-like plaque deposition, TMEM97 (sigma-2) modulators improved cognitive deficits.[25, 73] In human cases, we observed TMEM97 is increased in biochemically isolated synapses of Alzheimer’s patients compared to age-matched controls, using an unbiased proteomic approach.[24] Those findings and the fact that TMEM97 modulators are pharmacologically well studied, have brought the use of TMEM97 modulators into Phase II clinical trials for Alzheimer’s treatment.[11, 20]

However, the relationship between Aβ and TMEM97 in human cases was not previously clear. Our current results support a mechanistic explanation that includes a potentially direct interaction between Aβ and TMEM97, as suggested by the FRET findings (**Fig. 6**). Importantly, we found that this potential interaction may occur at the synapses, believed to be the earliest affected structure in the context of Alzheimer’s and the best pathological correlate of the characteristic cognitive decline.[16, 56, 65] Taken together, these findings link a therapeutic target (TMEM97) with a suspected pathological peptide (Aβ) in a key cellular structure (the synapse).

**Figure 6.**
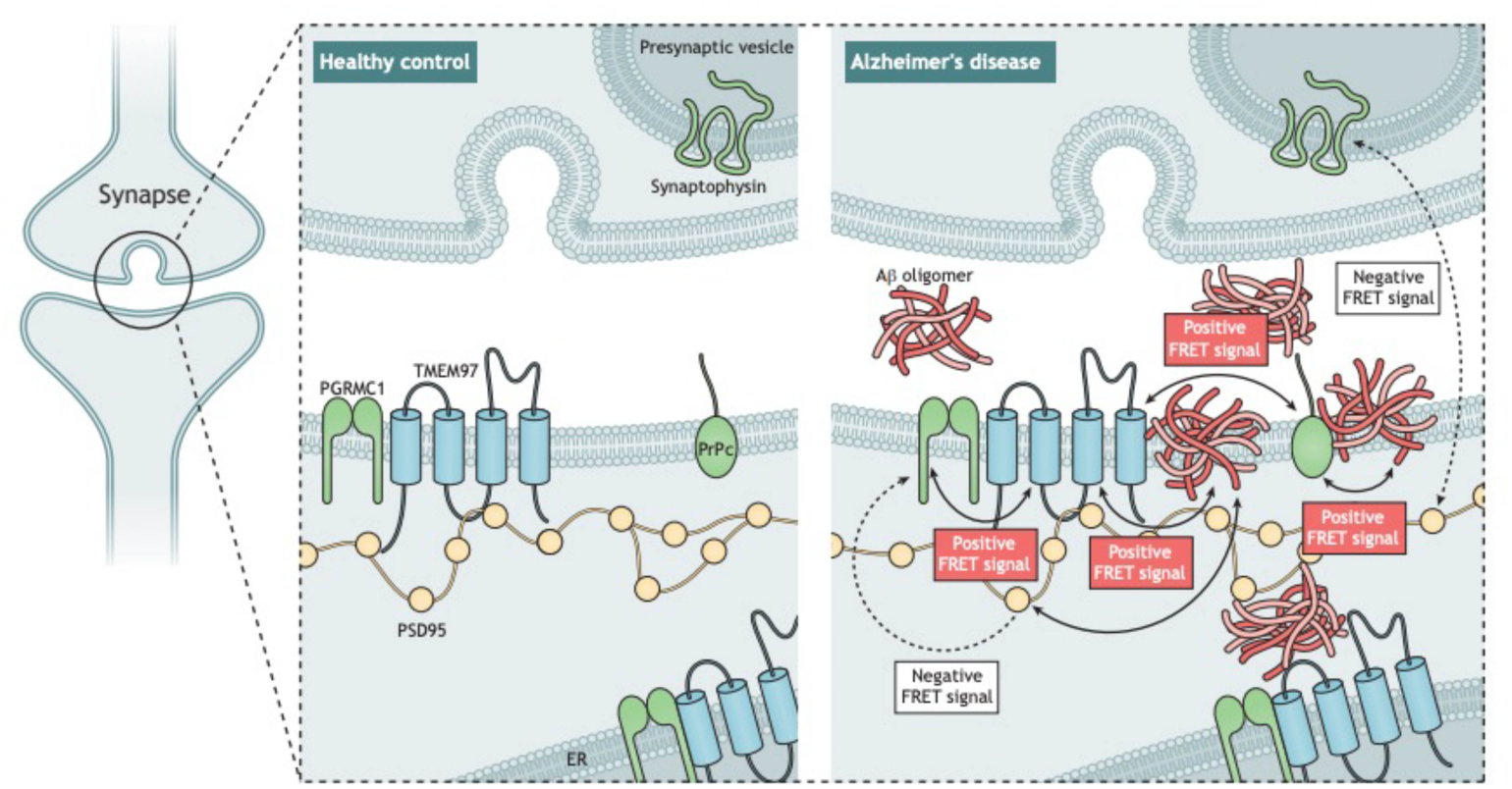
Model of synaptic interactions of Aβ. Based on our study, we observe that Aβ is in close proximity to TMEM97, PSD95, and PrPc. Also, TMEM97-PGRMC1 and TMEM97-PrPc were found close enough to generate FRET signals. There was no FRET signal generated between PGRMC1 and PSD95 nor between PSD95 and synaptophysin which should not be in close enough proximity to generate a signal. These data are consistent with Aβ being a binding partner of these synaptic proteins either at the synaptic membrane or potentially within the post-synapse at spine apparatus.

The relationship between Aβ and synapses has been widely studied.[18] It has been shown that Aβ can be found in synapses of Alzheimer’s cases,[6, 34, 51] but the mechanisms by which Aβ induces synaptic toxicity remain unclear. The study of synaptic binding partners of Aβ yielded many candidates – reviewed in.[30, 46, 63] The most studied binding partner is the cellular prion protein (PrP_c_), which through a cascade involving a complex with mGluR5 may lead to toxicity independently or via tau.[37, 62, 68, 75] Other binding partners that have been suggested to bind Aβ include the α7-nicotinic receptor,[49] Ephrin A4 (EphA4),[69] PSD95[35, 50, 52] or LilrB2[32]. It is likely that Aβ is in fact interacting with more than one protein.[62] Due its hydrophobic nature, Aβ binds to lipid membranes interrupting membrane fluidity and destabilizing several membrane receptors.[38] It is relevant to note that most binding partners have not been studied in human brain nor using human derived Aβ species.[38, 63] Further, while Aβ fibrils bind non-specifically to a variety of surfaces, and Aβ monomers bind to several receptors when applied exogenously,[5] Aβ oligomers have been shown to bind saturably to a single site[25, 62] suggesting specific pharmacological interactions with receptors. Therefore, it has been tricky to determine which Aβ binding partners are relevant in living human brain. The structural state of Aβ (monomer, oligomer or fibril) recognized in the present array tomography studies is not clear, which is a limitation of the study, but within the limitations of the technique, we are able to observe proximity of synaptic proteins to a degree that has not previously been possible within human synapses.

Here we observe that in human synapses, TMEM97 and Aβ are close enough to generate a FRET effect, an observation that allows us to define close proximity, but does not conclusively show a direct interaction. The fact that we also found a FRET effect between Aβ and PrP_c_ in the same cases is in line with previous observations,[37, 62] and reinforces the idea of multiple synaptic binding partners of Aβ at the synapses of Alzheimer’s cases. This is important for the field as both modulating TMEM97 with CT1812 and modulating PrPc rescue cognition in plaque-bearing transgenic mice,[14, 25] and these data indicate their beneficial effects could be through a similar mechanism. Given that TMEM97 and PrPc and PrPc and Aβ interact, and given that PrPc has been well characterized to bind Aβ, while it is possible that the interaction with TMEM97 and Aβ is direct, it is also possible that the FRET signal between TMEM97 and Aβ is through a direct interaction of TMEM97 with PrPc, thereby enabling a FRET signal with TMEM97 and Aβ, which is bound to PrPc. With the current methodology, we cannot conclusively determine the proportion of Aβ bound to each potential receptor in synapses but our data do support the idea of protein complexes interacting with Aβ.

How this interaction may be leading to synaptic dysfunction and subsequent neurodegeneration is less clear. Several mechanisms have been proposed linking Aβ and synaptic dysfunction involving excitatory imbalance. [10, 38] The fact that we found increased levels of TMEM97 in human Alzheimer’s cases and a close proximity between TMEM97 and Aβ at the synapses led us hypothesise that TMEM97 may be involved in the pathogenesis of Alzheimer’s and synaptic dysfunction. In the Alzheimer’s mouse model included in this study we were able to see a reduction of the Aβ potential binding to TMEM97 – as reflected by the decrease of FRET signal – associated with increased estimated receptor occupancy of the investigational drug CT1812, only when using the 80% receptor occupancy threshold for which has been previously reported to be required for efficacy.

Given that variable levels of CT1812, both under and over the 80% RO threshold, were observed in this study, it is not surprising that there was no statistically significant change in mean densities of Aβ, TMEM97 or PSD95 or the synaptic localization of Aβ and/or TMEM97 in the CT1812 treated group compared to vehicle. Post-hoc analyses however focusing on changes that might be mediated in animals for which >80% RO was achieved was in support of efficacy studies which found this threshold must be achieved for efficacy to be observed: that the increase of estimated receptor occupancy by the drug significantly correlated with a decrease of synaptic FRET signal between Aβ and TMEM97 (*rho*=-0.94, *p=*0.017, **Fig. 4F**). The inverse correlation of increasing CT1812 levels with decreased interaction between Aβ and TMEM97 is consistent with the mechanism of action ascribed preclinically, showing a reduction in oligomeric Aβ binding to neuronal synapses in the presence of Sigma-2 receptor modulators,[27] and is supportive that CT1812 dosed orally can engage the target, the TMEM97 (sigma-2) receptor, in the brain, in a drug exposure-dependent manner.

Notably, only five treated mice exhibited drug concentrations above the 80% estimated receptor occupancy, the drug concentration threshold previously defined as effective.[25] The low number of mice with high levels of drug make us cautious about the correlation found while highlight an unexpected finding: a statistically significant increase in drug concentration in male when compared with female mice. None of the variables controlled in the present study could explain the drug concentration differences between males and females and therefore we hypothesise that sex-related biological differences in mouse may be underlying the drug metabolism or blood brain barrier penetration. This is the first study in which CT1812 has been administered to animal models in food, however human clinical trials suggest no difference in CT1812 pharmacokinetics in a fed or fasted state.[20] It is important to note that Izzo and colleagues found an improvement of cognition in mice exhibiting more than 80% estimated receptor occupancy, something only seen in one female of our study.[25] Further, Izzo and colleagues included only male mice in the study, and therefore the present findings on female mice should be taken into consideration to ensure the efficacy of treatments in future studies. These findings may be in line with the increasing body of literature describing sex differences in mice models of Alzheimer’s that may or may not be translated to human cases.[1, 2, 15]

Although we could see a reduction in the synaptic FRET signal of Aβ with TMEM97 in the >80% estimated receptor occupancy group, and a significant correlation with drug brain concentration in the treated group, the treatment of the Alzheimer’s mouse model with TMEM97 modulators did not result in a recovery of synaptic densities nor a decrease of Aβ synaptic localization. CT1812 has been previously demonstrated to selectively displace Aβ oligomers, but not monomer, from synaptic receptor sites and facilitate its clearance out of the brain,[28] suggesting that the Aβ that is interacting with TMEM97 observed in this study may be predominantly fibril as opposed to oligomeric. However, disrupting this interaction between TMEM97 and Aβ may be sufficient to improve synaptic function, without requiring a detectable change in synapse density, which could explain the behavioural recovery seen in previous mouse studies with CT1812 treatment.[25] Alternatively, the 28 day treatment period used here may not have been sufficiently long to observe a change in synaptic density; previous studies demonstrating CT1812-mediated improvement in cognitive performance were conducted following 9-10 weeks of administration.[28] Lastly, whereas data herein were generated in the APP/PS1+Tau mouse model, behavioural data was generated in the Swedish London APP model and it’s possible the forms of Aβ most abundant or relative ratios of oligomeric and fibril forms of Aβ are different across Alzheimer’s models.[28]

A previously published model of CT1812’s mechanism of action proposes that the Sigma-2 receptor complex regulates other Aβ oligomer receptors (composed of LilRB2, NGR and PrPc), and when CT1812 binds to TMEM97, allosteric interactions between the Sigma-2 receptor and the oligomer receptor change the oligomer receptors’ shape, destabilize the binding pocket, and increase the off-rate of Aβ oligomers from their receptor. Therefore suggesting that CT1812 does not compete directly with oligomers at the same site.[28] In tumor cells, the canonical Sigma-2 ligand DTG binds to Sigma-2 receptors at a location on the TMEM97 protein,[3] and CT1812 displaces radiolabeled DTG binding, but the precise binding location of CT1812 has not been directly determined. Our data indicate that TMEM97 and Aβ are in close proximity where they could be binding, but we cannot rule out that Aβ may be binding to other nearby proteins instead of directly interacting with TMEM97.

In conjunction with FRET showing a synaptic localization, transcriptomic data from human iPSC derived neuronal cultures treated with soluble fraction of Alzheimer’s brain homogenate with the Sigma-2 modulator CT1812 support a synaptic localization and mechanism. UniProt pathway analysis identified “Synapse” as the most significant term in transcripts altered by CT1812 and the top gene ontology terms important are synaptic and include postsynaptic density and postsynaptic membrane. Of the eight transcripts regulated by CT1812 using a highly stringent statistically criterion (B-H adjusted p<0.05) was Protocadherin gamma-B4 (PCDHGB4), a cell adhesion molecule on neuronal synapses, which is involved in synaptic maturation and stabilization.[36]

Other amongst the eight top-most significant transcripts altered suggest a role for CT1812 in modulating glia. In the cultures studied, ∼16% of cells were GFAP+ astrocytes and data therefore represents a context of cortical neuron and astrocyte interaction. One of the transcripts significantly changed by CT1812 treatment, CHI3L1, which encodes YKL-40 protein, is expressed in astrocytes and is a biomarker of inflammation in Alzheimer’s disease.[48] Beyond the transcript level view, pathway analysis using MetaCore (unadjusted *p*-value<0.05) revealed enrichment of astrocytic and immune/inflammatory pathways with Aβ treatment compared to immunodepleted Aβ treatment, including the JAK/STAT pathway known to be involved in astrocyte differentiation and function, indicated by Leukemia Inhibitory Factor (LIF) and Leukemia Inhibitory Factor Receptor (LIFR) enrichment, a known pro-inflammatory cytokine involved in the differentiation of neuronal precursor cells into astrocytes (**Fig. 5F**). We also see in mice that treatment with CT1812 reduces astrogliosis in hippocampus, further supporting an important role for astrocytes in protecting synapses and ultimately cognition. Together, data suggest a role for CT1812 in modulating inflammatory pathways, perhaps indirectly by glia that may detect changes in synapse health or activity.

Regarding the mechanism by which TMEM97-Aβ interaction may be linked to synaptic dysfunction, Also in line with the hypothesis of Aβ binding to lipid membranes,[38] TMEM97 is thought to be an endo-lysosome-related protein which has itself been linked to cellular toxicity and is essential for the internalization of cholesterol molecules like LDL through the formation of a complex with PGRMC1.[4, 55] Our transcriptomic data from human iPSC derived neurons challenged with Alzheimer’s brain derived Aβ confirms neuroinflammatory and neurodegenerative related changes with Aβ challenge. CT1812 treatment of these cells caused multiple changes in transcripts involved in synaptic function, consistent with the ability of CT1812 to influence synapses and potentially ameliorate damage in Alzheimer’s.

In summary, in human Alzheimer’s brains we found increased synaptic levels of TMEM97 and close colocalization of TMEM97 with Aβ in synapses. This supports the idea that TMEM97 is a synaptic binding partner for Aβ either directly or indirectly, which is important as this interaction can be modulated by drugs.

## Abbreviations

Aβ: amyloid beta
TMEM97: transmembrane protein 97
PSD95: postsynaptic density 95
PrPc: cellular prion protein
PGRMC1: Progesterone receptor membrane component 1.

## Acknowledgements

The authors thank our brain tissue donors and their families for their generous donations. Authors gratefully acknowledge membership of Edinburgh Neuroscience. Some of the control participants in the human study were from the Lothian Birth Cohort 1936, thus we wish to thank the cohort and research team supported by Age UK (Disconnected Mind project) in The University of Edinburgh Centre for Cognitive Ageing and Cognitive Epidemiology, funded by the Biotechnology and Biological Sciences Research Council (BBSRC) and Medical Research Council (MRC) ((MR/K026992/1). We also acknowledge Neil Smith for artwork in Figure 6.

## Funding

This work was supported by Alzheimer’s Society (project grant AS-PG-15b-023), Alzheimer’s Research UK, the European Research Council (ERC) under the European Union’s Horizon 2020 research and innovation programme (Grant agreement No. 681181), the University of Edinburgh (Chancellor’s Fellow start-up funding), Wellcome Trust Institutional Strategic Support Fund, the UK Dementia Research Institute which receives its funding from DRI Ltd., funded by the UK Medical Research Council, Alzheimer’s Society, and Alzheimer’s Research UK (UKDRI-Edin0005), and BBSRC Institute Strategic Programme funding.

## Competing interests

TSJ is a member of the Scientific Advisory Board of Cognition Therapeutics. NI, LW, and SC were employees of Cognition Therapeutics, and MEH and AOC are current employees of Cognition Therapeutics.

## Supplementary material

Supplementary material is available at *Brain* online.

## Notes

### Competing Interest Statement

TSJ is a member of the scientific Advisory Board of Cognition Therapeutics. LW, AOC, MH, NI and SC are or were employees of Cognition Therapeutics.

### Summary of Updates

New iPSC data has been added after peer review.

