## Supplementary Material for "Transmembrane protein 97 is a potential synaptic amyloid beta receptor in human Alzheimer’s disease"

### TMEM97 increases in synapses and is a potential synaptic Aβ binding partner in human Alzheimer's disease

**Table S1. Mice included in array tomography-FRET experiments.**

**Table S2. Antibodies.**

**Fig. S1. Image analysis pipeline.**

**Fig. S2. Effect of sigma-2 receptor antagonist on PSD95, Aβ and tau in the non-transgenic control mice and APP/PS1+Tau mice.**

**Fig. S3. Plaque and astrocyte burdens in transgenic mice treated with CT1812 or vehicle.**

**Table S1. Mice included in array tomography-FRET experiments.**

| Case | Animal ID | Treatment | Age at death | Sex | Brain CT1812 (ng/mL) | Blood CT1812 (ng/mL) | Estimated receptor occupancy (%) |
|---|---|---|---|---|---|---|---|
| 1 | G353 | vehicle | 10,2 | f | 0 | NA | 0 |
| 2 | G394 | vehicle | 10,1 | f | 0 | 0 | 0 |
| 3 | G395 | vehicle | 10,1 | f | 0 | 0 | 0 |
| 4 | G439 | vehicle | 9,9 | f | 0 | 0 | 0 |
| 5 | G443 | vehicle | 9,9 | m | 0 | 0 | 0 |
| 6 | G454 | vehicle | 9,9 | m | 0 | 0 | 0 |
| 7 | G455 | vehicle | 9,9 | m | 0 | 0 | 0 |
| 8 | G458 | vehicle | 9,9 | m | 0 | 0 | 0 |
| 9 | G362 | CT1812 | 10,1 | f | 9,7 | 0,9 | 72,5 |
| 10 | G365 | CT1812 | 10,1 | f | 9,0 | 0,7 | 71,0 |
| 11 | G372 | CT1812 | 9,9 | m | 12,3 | 0,9 | 77,0 |
| 12 | G413 | CT1812 | 10,1 | f | 18,8 | 1,1 | 83,7 |
| 13 | G421 | CT1812 | 10,0 | m | 22,0 | 0,1 | 85,7 |
| 14 | G429 | CT1812 | 9,9 | m | 47,5 | 7,1 | 92,8 |
| 15 | G435 | CT1812 | 9,9 | m | 15,4 | NA | 80,7 |
| 16 | G444 | CT1812 | 9,9 | m | 30,9 | 1,9 | 89,4 |
| 17 | G519 | CT1812 | 9,9 | f | 7,9 | 0,5 | 68,4 |
| 18 | G520 | CT1812 | 9,9 | f | 4,1 | 0,2 | 52,9 |

Abbreviations: f, female; m, male; NA, not available.

**Table S2. Antibodies.**

| Primary Antibodies | Host specie | Reactivity specie | Supplier | Catalogue # | Dilution | Study |
|---|---|---|---|---|---|---|
| PSD95 | Guinea Pig | Human | Synaptic Systems | 124-014 | 1:50 | AT-FRET main study |
| Aβ (6E10) | Mouse | Human | Biolegend | 39320 | 1:200 | AT-FRET main study |
| TMEM97 | Rabbit | Human | Novus Biologicals | NBP1-30436 | 1:100 | AT-FRET main study |
| PrPc (EP1802Y) | Rabbit | Human | Abcam | ab52604 | 1:50 | AT- FRET other interactors |
| Synaptophysin (Sy38) | Mouse | Human | Abcam | Ab8049 | 1:50 | AT-FRET other interactors |
| PSD95 | Rabbit | Human | Cell Signaling | D27E11 | 1:50 | AT-FRET other interactors |
| PGRMC1 | Goat | Human | Abcam | Ab48012 | 1:50 | AT-FRET other interactors |

| Secondary Antibodies | Host specie | Reactivity specie | Supplier | Catalogue # | Dilution | Study |
|---|---|---|---|---|---|---|
| Alexa Fluor 488® | Donkey | Guinea Pig | Jackson Immuno | 706-545-148 | 1:50 | AT-FRET main study |
| Cy™3 | Donkey | Mouse | Jackson Immuno | 715-165-150 | 1:50 | AT-FRET main study |
| Cy™5 | Donkey | Rabbit | Jackson Immuno | 711-175-152 | 1:50 | AT-FRET main study |
| Cy™3 | Goat | Mouse | Jackson Immuno | 115-165-146 | 1:50 | AT-FRET positive control |
| Cy™5 | Donkey | Goat | Jackson Immuno | 705-175-147 | 1:50 | AT-FRET positive control |

**Fig. S1. Image analysis pipeline.**

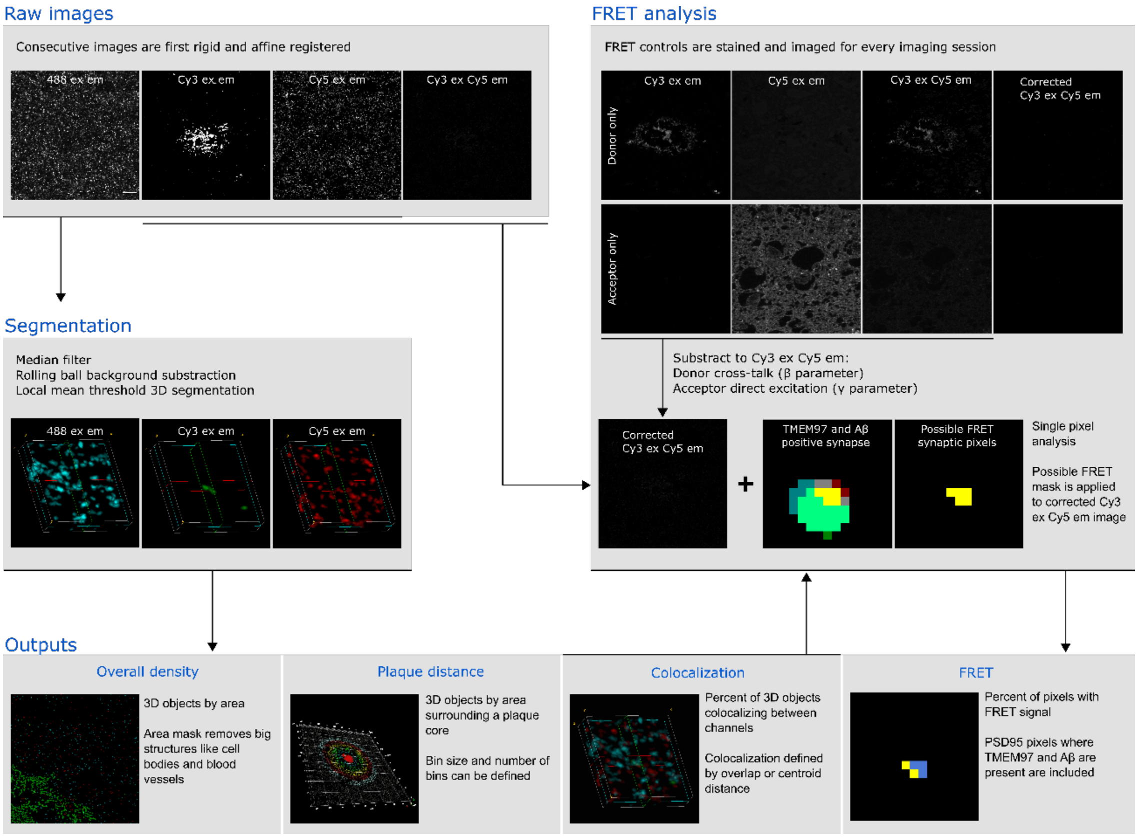


The diagram describes the basic image analysis steps performed in the current study. All the processes were combined into an in-house algorithm available at: *add upon acceptance for publication*.

**Fig. S2. Effect of sigma-2 receptor antagonist on PSD95, Aβ and tau in the non-transgenic control mice and APP/PS1+Tau mice.**

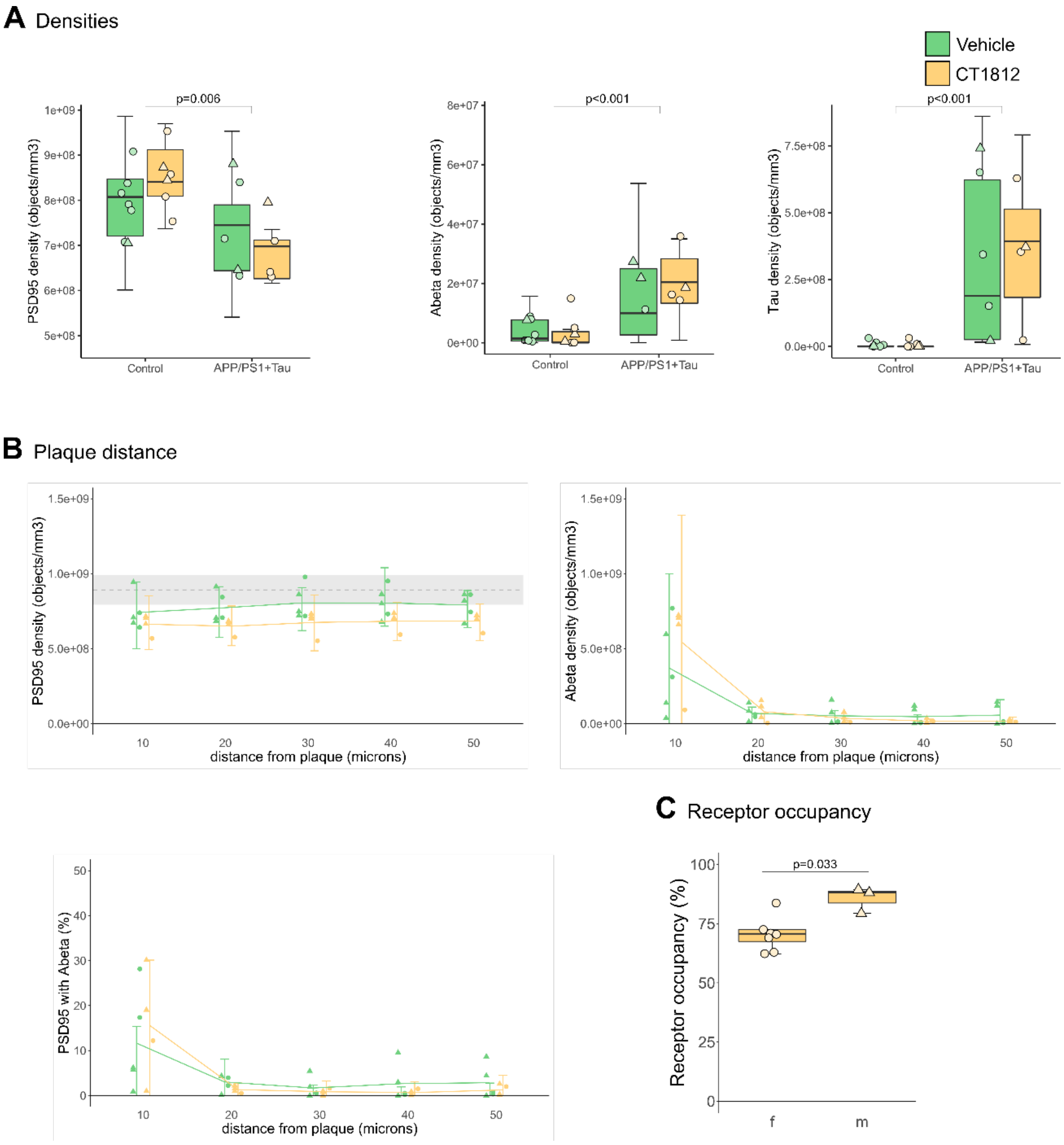


In **A** are quantified the densities of object found for either PSD95, Aβ or tau in the non-transgenic control mice and the APP/PS1+Tau mice model. In **B**, plaque distance density of either PSD95, Aβ, or PSD95 that contain Aβ in the APP/PS1+Tau are plotted. Grey dotted line show the mean of control mice and the SD is shown in grey. In **C**, is plotted the estimated receptor occupancy by the drug in the treated mice, independently of the genotype. Boxplots show quartiles and medians calculated from each image stack. Data points refer to case means (females, triangles; males, circles). Analysis with linear mixed effects models including treatment group and sex interaction. Abbreviations: f, female; m, male.

**Fig. S3. Plaque and astrocyte burdens in transgenic mice treated with CT1812 or vehicle.**

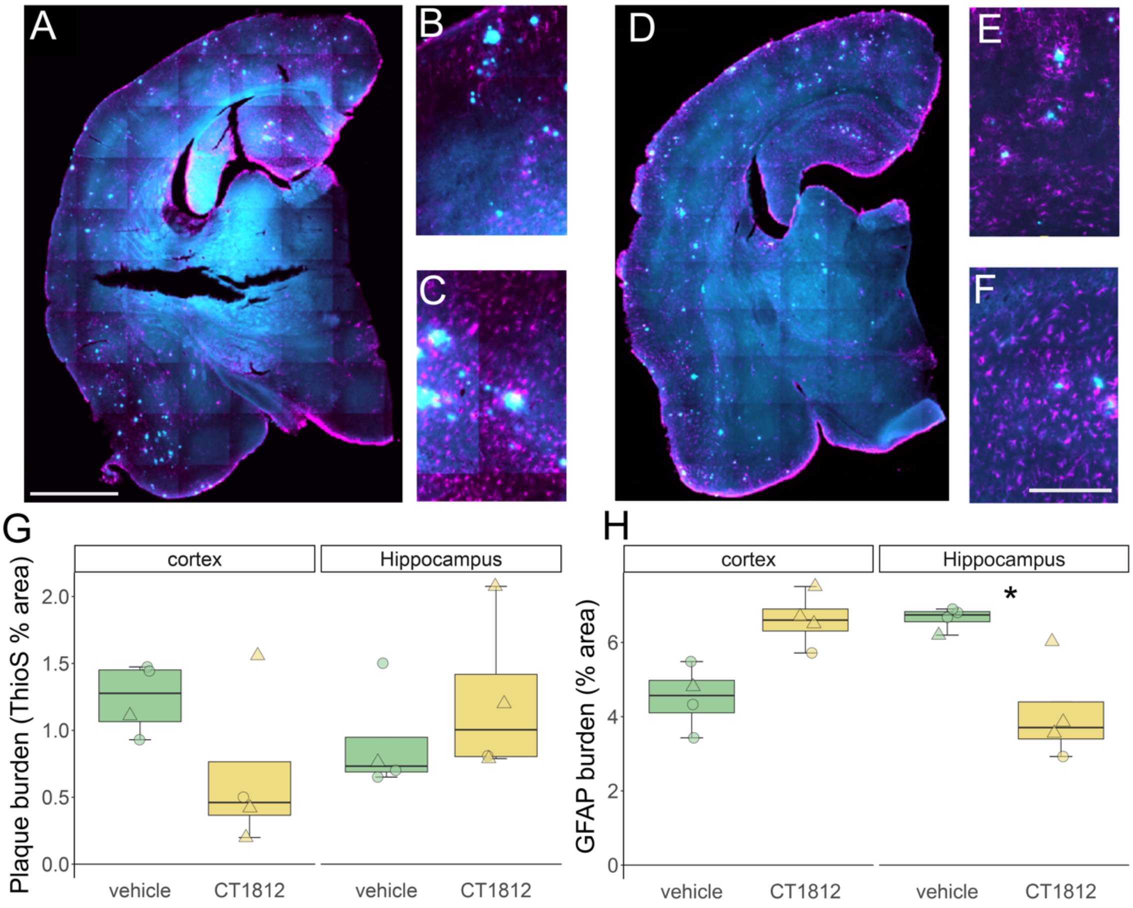


Thioflavin S plaque burden and GFAP astrocyte burden were examined on 50 micrometer floating sections (3 per mouse) in animals treated with vehicle (A-C) or CT1812 (only those with >80% estimated receptor occupancy included in the study, representative images D-F). Tilescans of the entire section were taken (A, D) and analysis carried out in cortex (B, E) and hippocampus (C,F). While there were no significant differences in plaque burdens (G), ANOVA on linear mixed effects models of astrocyte burdens shows a significant interaction between treatment and brain region ($F[1,16]=9.46$, $p=0.007$) and a significant post-hoc difference decrease in astrocyte burden in hippocampus with CT1812 treatment ($t=2.43$, $p=0.03$). Scale bars represent 1mm in A and D and 250 μm in B, C, E, and F.
